# Integrating current analyses of the breast cancer microbiome

**DOI:** 10.1101/2022.10.02.510553

**Authors:** Sidra Sohail, Michael B. Burns

## Abstract

Breast cancer is the second leading cause of cancer death for women in the US (*American Cancer Society: About Breast Cancer*, n.d.). Many cancer types have significant associations with their resident microbial communities - emerging evidence suggests that breast cancers also interact with the local tissue-associated microbiota. Studies have examined the relationship between breast cancer and its microbiome, however, the studies varied in their approaches used to evaluate these relationships. Microbiome research advances rapidly and analysis pipelines and databases are updated frequently. This dynamic environment makes inter-study comparisons and superficial evaluations challenging as no two studies are using the same standards for evaluation.

Researchers have observed the microbiota of tumor tissue, surrounding normal sites, and healthy breast tissue from non-cancer individuals (Hieken et al., 2016; Urbaniak et al., 2016; Xuan et al., 2014), but they have not been able to translate their findings into information that can be used for breast cancer treatment or detection nor address what affect studying different variable regions has in their analysis. Within the majority of these studies, comparisons of the tumor tissue with adjacent normal tissue has revealed differences. This project will integrate all available studies related to breast cancer and the mammary microbiome to 1 reassess the original findings in light of advances in this rapidly progressing field and 2 incorporate all the data available as a large meta-analysis to identify general trends and specific differences across patient cohorts and studies.

## Introduction

Breast cancer is the second leading cause of cancer death for women in the US (*American Cancer Society: About Breast Cancer*, n.d.). Many cancer types have significant associations with their resident microbial communities - emerging evidence suggests that breast cancers also interact with the local tissue-associated microbiota (Chan et al., 2016; Hieken et al., 2016; Urbaniak et al., 2016; Xuan et al., 2014). Studies have examined the relationship between breast cancer and its microbiome, however, the studies varied in their approaches used to evaluate these relationships. Microbiome research advances rapidly and analysis pipelines and databases are updated frequently. This dynamic environment makes inter-study comparisons and superficial evaluations challenging as no two studies are using the same standards for evaluation.

Researchers have observed the microbiota of tumor tissue, surrounding normal sites, and healthy breast tissue from non-cancer individuals (Hieken et al., 2016; Urbaniak et al., 2016; Xuan et al., 2014), but they have not been able to translate their findings into information that can be used for breast cancer treatment or detection nor address what affect studying different variable regions has in their analysis. Within the majority of these studies, comparisons of the tumor tissue with adjacent normal tissue has revealed differences. The microbiota differ drastically with the malignant tissue showing an increased abundance of pro-inflammatory genera and a decrease in bacterial community diversity and bacterial load (Hieken et al., 2016; Xuan et al., 2014). The depleted bacterial diversity in the malignant tissue can be potentially explained by the hypoxic, inflammatory microenvironment of tumor tissue. There is a decrease in the bacterial load in advanced tumors, and also a reduction in the antibacterial response in the breast tumor tissue where more severe tumors have a lower abundance of innate immune receptors in breast tissue (Xuan et al., 2014). However, when evaluating the microbiota of other cancer types, the cancers harbor more diverse communities (Burns et al., 2015; Mira-Pascual et al., 2015).

In comparisons of healthy and malignant breast tissue (Hieken et al., 2016), *Proteobacteria* and *Firmicutes* show increased abundance in tumor tissue (Urbaniak et al., 2016; Xuan et al., 2014). **However, there is not a functional or clear mechanistic explanation of these differences nor any inkling of how this translates to potential treatment or improvements in diagnosis for breast cancer**. Also, each study uses different bioinformatic methods, data formats, and variable regions further blurring the applicability of the results of each study to our understanding of the disease generally. The available studies (Chan et al., 2016; Hieken et al., 2016; Urbaniak et al., 2016) fail to address the question of what the microbial composition in the mammary microbiome entails for mammary health, and whether looking at the different variable regions in their 16S rRNA analysis has any major effects on their results. It is necessary to combine the findings from these studies with respect to the different variable regions and patient cohorts. There are a variety of taxonomic databases and each study has used a different database. Databases have greatly evolved overtime of which the Greengenes database (McDonald et al., 2012) is deprecated, and some of the current databases are RDP (Cole et al., 2005) and SILVA (Pruesse et al., 2007). However, prior versions of these databases are incomplete relative to the newer, updated ones. The microbiome field is rapidly changing as there are different variable regions, different databases, and different pipelines available. Therefore, in this work, we are performing a retrospective analysis of existing studies on the breast cancer microbiome.

## Methods

### Dataset overview

The study *The Microbiome of Aseptically Collected Human Breast Tissue in Benign and Malignant Disease* by Hieken et al. in August 2016, targets the V3-V5 variable region of their 16S rRNA data and compares the microbial communities between malignant and benign tumors of patients with breast cancer, using normal adjacent tissues as controls. Their data is comprised of 98 forward and 98 reverse fastq files with a patient sample size of 33 patients **(Fig. 1A)**, of which there were 16 women with benign disease and 17 women with malignant disease. The tissues of interest are buccal swab, skin swab, breast tissue, and breast skin tissue, which is also referred to as skin tissue in the paper. The sequencing data for this study is available from the NCBI SRA (*Sequence Read Archive (SRA)*, n.d.) and was downloaded from the SRA with accession number PRJNA335375, using the SRA explore interface at sra-explorer.info (Ewels et al., n.d.). They analyzed their 16S rRNA paired-end data by implementing the IM-TORNADO bioinformatics pipeline (Jeraldo, 2016/2020) using the Greengenes version 13.5 reference database (McDonald et al., 2012). Furthermore, they implemented two alpha diversity measures which were the observed OTU number and the Shannon Index, two beta diversity measures which were the unweighted and weighted UniFrac distances (Lozupone & Knight, 2005), and performed differential abundance analysis to identify significant taxa. The diversity measures were calculated using an OTU table and phylogenetic tree. The observed OTU number reflects the species richness and the Shannon index reflects the species evenness. The unweighted UniFrac (Lozupone et al., 2011) measures differences in community presence such as whether or not there is an OTU present, and the weighted UniFrac measures differences in community abundance. This paper did not perform multiple correction and reported unadjusted p-values.

**Figure 1.**
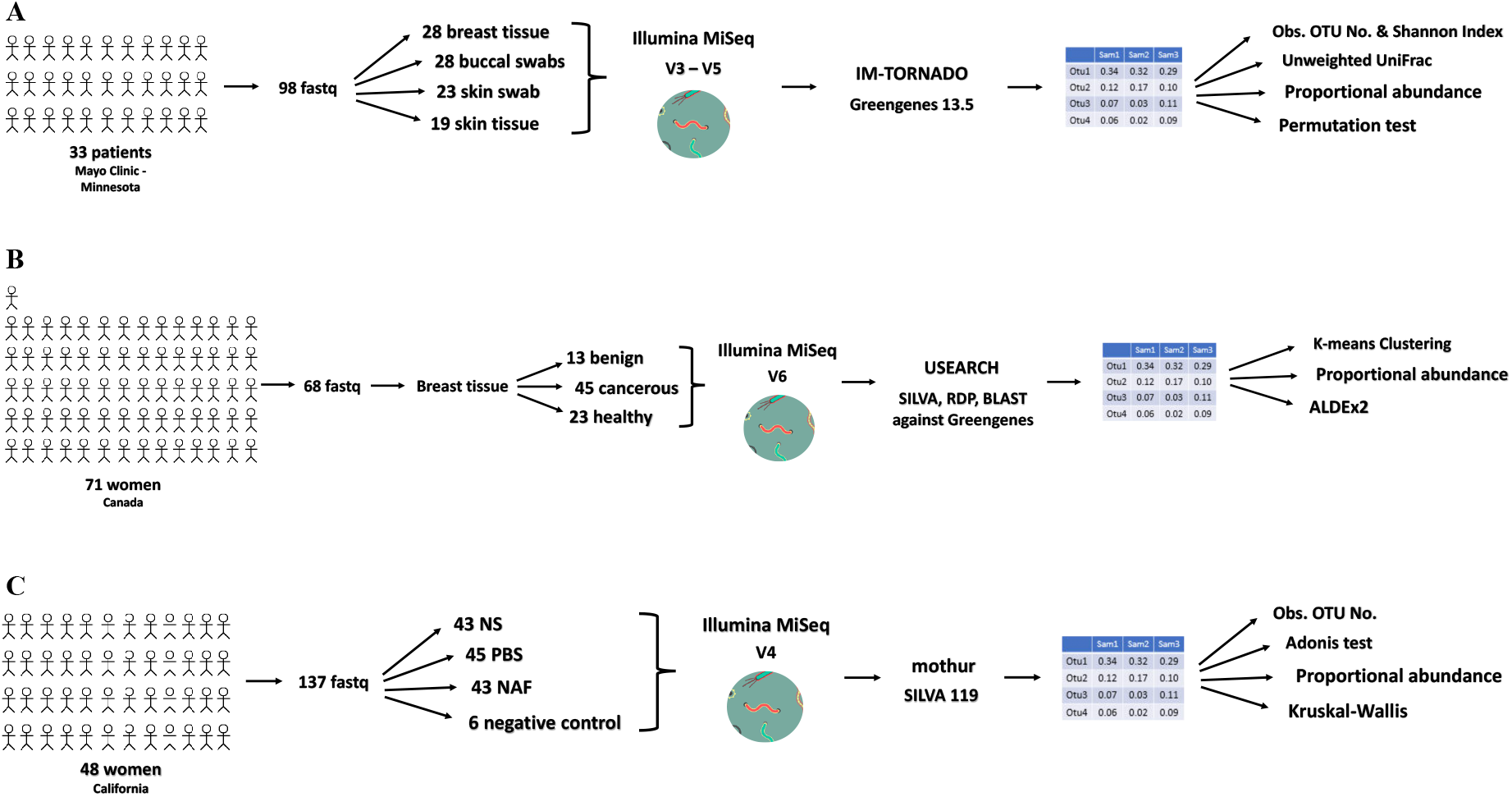
Overview of the studies of interest. (A) Hieken study overview outlining their patient sample size, number of samples, sequencing technology and variable region, bioinformatics pipeline and taxonomic reference database, and downstream analyses. (B) Urbaniak study overview outlining their patient sample size, number of samples, sequencing technology and variable region, bioinformatics pipeline and taxonomic reference database, and downstream analyses. (C) Chan study overview outlining their patient sample size, number of samples, sequencing technology and variable region, bioinformatics pipeline and taxonomic reference database, and downstream analyses.

The study *The Microbiota of Breast Tissue and Its Association with Breast Cancer* by Urbaniak et al. in August 2016, targets the V6 variable region of their 16S rRNA data and compares the microbial composition between breast tissue from healthy control women and normal adjacent breast tissue from women with breast cancer, and they also compare normal adjacent breast tissue and breast tumor tissue. Their data is comprised of 68 merged fastq files with a patient sample size of 71 patients **(Fig. 1B)**, of which there were 13 benign, 45 cancerous, and 23 healthy samples. The samples of interest in this study are malignant, benign, and healthy samples from breast tissue in this paper, where the samples are normal adjacent tissue from women with breast cancer which is either benign or malignant and tissue from healthy controls. The sequencing data for this study is available from the NCBI SRA (*Sequence Read Archive (SRA)*, n.d.) and was downloaded from the SRA with accession number SRP076038, using the SRA explore interface at sra-explorer.info (Ewels et al., n.d.). They analyzed their 16S rRNA paired-end data by clustering the reads into OTUs by 97% identity using the Uclust algorithm within USEARCH version 7 (Edgar, 2010). Taxonomic assignment of each OTU was made by extracting best hits from SILVA database (Pruesse et al., 2007), manually verifying using Ribosomal Database Project (RDP) SeqMatch tool (Cole et al., 2005), and then using BLAST (Altschul et al., 1990) against the Greengenes database (McDonald et al., 2012), where the hits with the highest percent identity and coverage were used to assign taxonomy. The OTU sequences were aligned using MUSCLE (Edgar, 2004) and were inputted to FASTTREE (Price et al., 2009) to build a tree of OTU sequences where PCoA plots of weighted UniFrac distances (Lozupone et al., 2011) were made in QIIME (Caporaso et al., 2010) using this tree of OTU sequences.

Additionally, unsupervised k-means clustering was performed using euclidean distances of center-log ratio (CLR) transformed data and used the ALDEx2 R package (Gloor et al., 2022) to measure the relative abundances of genera where Benjamini-Hochberg corrected p-value of the Wilcoxon rank test was used to test for significance. Furthermore, to test for differences between microbiota a microbiome regression-based kernel association test (mirkat) was performed in R using the MiRKAT package (Zhao et al., 2015) where a kernel metric was built using UniFrac distances and Bray-Curtis dissimilarity metric. UniFrac distances and Bray-Curtis dissimilarity metric were both used in the kernel metric simultaneously.

The study *Characterization of the microbiome of nipple aspirate fluid of breast cancer survivors* by Chan et al. in June 2016, targets the V4 variable region of their 16S rRNA data and investigates the microbial community present in nipple aspirate fluid (NAF) and their potential association with breast cancer by comparing NAF between healthy control samples and women with history of breast cancer samples. Their data is comprised of 137 forward and 137 reverse fastq files with a patient sample size of 48 patients **(Fig. 1C)**, of which there were 23 healthy control women and 25 women with a history of cancer samples. There are healthy control and women with a history of breast cancer samples and tissues of interest are NAF, nipple skin (NS), and post-betadine skin (PBS) in this paper. The women with a history of breast cancer are also referred to as cancer samples in this section. The sequencing data for this study is available from the NCBI SRA (*Sequence Read Archive (SRA)*, n.d.) and was downloaded from the SRA with accession number PRJNA314877, using the SRA explore interface at sra-explorer.info (Ewels et al., n.d.). There were OTUs detected in empty control Eppendorf tubes and were removed from analyses to account for contaminating microbial 16S rDNA sequences from extraction kits and reagents. They analyzed their 16S rRNA paired-end data by implementing the Schloss MiSeq standard operating procedure in mothur (Schloss et al., 2009) using the SILVA version 119 reference database (Pruesse et al., 2007). They implemented an alpha diversity measure which was the observed OTU number, a beta diversity measure which was the analysis of variance (Adonis) test using Bray-Curtis distance and PCoA, and performed differential abundance analysis through the Kruskal-Wallis test to identify significant taxa. The Adonis test was used to measure community composition differences and a nonparametric Kruskal-Wallis test was used to test whether the OTU relative abundances were statistically significant between the healthy control and breast cancer samples for NAF, NS, and PBS samples. The OTU table was rarefied for calculating dissimilarity measures and PCoA to account for any bias from uneven sequencing depth. The statistical analyses used a p-value cutoff of 0.05 and false discovery correction was not applied when comparing the OTU relative abundances; however, a multiple test correction was applied when performing functional prediction with PICRUST (Langille et al., 2013) and KEGG (Kanehisa & Goto, 2000).

### Preprocessing

In the Hieken and Chan studies, prior to beginning denoising, the primers used to amplify different regions of the 16S rRNA gene were trimmed from the unzipped paired end fastq files using cutadapt version 1.15 (Martin, 2011). In the Urbaniak study, upon downloading the fastq files from the SRA (*Sequence Read Archive (SRA)*, n.d.), the files were already merged and pre-processed; hence, primers were not trimmed.

The Hieken study targeted the V3-V5 regions, the Chan study targeted the V4 region, and the Urbaniak study targeted the V6 region of the 16S rRNA gene. In the Hieken study, the 357F primer used was AATGATACGGCGACCACCGAGATCTACACTATGGTAATTGTCCTACGGGAGGCAGC AG and the 926R primer used was CAAGCAGAAGACGGCATACGAGATGCCGCATTCGATXXXXXXXXXXXXCCGTCAAT TCMTTTRAGT. In the Chan study, the F515 primer used was AATGATACGGCGACCACCGAGACGTACGTACGGTGTGCCAGCMGCCGCGGTAA and the R806 primer used was CAAGCAGAAGACGGCATACGAGATXXXXXXXXXXXXACGTACGTACCGGATACHV GGGTWTCTAAT. In the Urbaniak study, the V6-forward primer used was 5′ACACTCTTTCCCTACACGACGCTCTTCCGATCTnnnn(8)CWACGCGARGAACCTTAC C3′ and the V6-reverse primer used was 5′CGGTCTCGGCATTCCTGCTGAACCGCTCTTCCGATCTnnnn(8)ACRACACGAGCTGAC GAC3′.

After trimming the primers, FastQC version 0.11.5 (*Babraham Bioinformatics - FastQC A Quality Control Tool for High Throughput Sequence Data*, n.d.) was used to assess the Phred quality score of each forward and reverse fastq file as well as to check for potential artifactual sequences in the dataset. In the Urbaniak study, the files were already merged and pre-processed; hence, FastQC was not used to assess the quality scores.

### DADA2 Pipeline Analysis

The trimmed reverse fastq files from the Hieken and Chan data were of very low quality and few of them made it through the quality control steps within the pipeline; therefore, only forward fastq files were included in the analysis, as these reads met our quality control standards. The trimmed forward fastq files from the Hieken and Chan data and merged fastq files from the Urbaniak data were analyzed using default parameters in the DADA2 1.16 pipeline (*DADA2 1*.*16 Pipeline*, n.d.), with changes in the filterAndTrim command, makeSequenceTable command, and the mergers command. In the filterAndTrim command, the reverse file input and truncLen parameter were removed and the maxEE value was adjusted from the default maxEE = c(2,2) to maxEE = 3. These changes were implemented to allow for flexibility and to relax the stringency in our analysis so to prevent massive removal of sample reads which would make interpretation of the results impossible. Also, the multithread parameter was set to equal two to avoid memory issues. In the makeSequenceTable command, the dadaFs variable was used to make the seqtab variable instead of the default mergers variable. The mergers command was removed as the analysis was performed with only the forward fastq files as input. Additionally, the SILVA version 138 reference database (Pruesse et al., 2007) compatible with DADA2 (Callahan et al., 2016) was downloaded from the Zenodo site (McLaren & Callahan, 2021). The tax_glom and transform_sample_counts functions in phyloseq were implemented to make the proportional abundance plots. Additionally, the microshades package was used to make colorblind friendly proportional abundance plots and the code can be found at the microshades website (*A Custom Color Palette for Improving Data Visualization*, n.d.). In order to make the microshades plot, the phyloseq object is used as input to the prep_mdf function. After making the proportional abundance plots, phylogenetic analyses were performed.

### Phylogenetic Analysis

The phylogenetic code was adapted from the compbiocore github site (*Computational Biology Core - Brown University*, n.d.), specifically commands from sequences<-getSequences(seqtab.nochim) up to dm <-dist.ml(phang.align) were implemented. The tree was generated using dm as input to the upgma command from the phangorn R package (Schliep et al., 2021). The ASV table and taxonomy table outputs from DADA2 along with the metadata and phylogenetic tree files were used as input to the MicrobiomeAnalyst web interface (Dhariwal et al., 2017) for visual exploratory data analysis.

### Statistical Analysis

In order to calculate the UniFrac distances for the Hieken and Urbaniak data (Lozupone & Knight, 2005), the GUniFrac function from the GUniFrac R package (Chen et al., 2022) was implemented where the input was the ASV table output from the DADA2 analyses. The UniFrac distances were used to perform beta diversity analyses for the Hieken and Urbaniak data.

Additionally, to perform statistical analyses for the Hieken data, a 10% prevalence filter and total sum scaling was applied to the ASV table to identify differentially abundant taxa. A permutation test was also performed for the Hieken data to identify significant ASVs of which were visualized through bar plots. In order to perform statistical analyses for the Urbaniak data, both UniFrac and Bray-Curtis distances were used to perform beta diversity analyses where only UniFrac distances were used to perform K-Means Clustering. Additionally, for the Urbaniak data, the ALDEx2 R package (Gloor et al., 2022) was used to measure the relative abundances of statistically significant genera. The ASVs in the ALDEx2 output were used to link associated taxa at the genus level to the ALDEx2 output and each genus in the output was visualized through box plots.

In order to perform statistical analyses for the Chan data, the Adonis test and Kruskal-Wallis test were run. The Adonis test was run to measure community composition differences and a nonparametric Kruskal-Wallis test was run to test whether relative abundances of ASVs were statistically significant between the healthy control and breast cancer samples for NAF, NS, and PBS samples. The p-values reported for the Adonis and Kruskal-Wallis tests are unadjusted p-values as the Chan study also reported unadjusted p-values. The Bray-Curtis distance and a proportionally rarefied phyloseq object were used to perform the Adonis test. The Kruskal-Wallis test was then run for each ASV from a rarefied phyloseq object, and ASVs that are not significant are filtered out. The significant ASVs were then linked with their associated taxa and the abundance values for each taxon were visualized through a dot plot.

## Results

### Hieken et al. Re-analysis Results

In the alpha diversity analysis, we have used the observed OTU number and Shannon Index to look at breast and skin tissue microbiota. The observed OTU number shows difference between the two tissues with an increased abundance of breast tissue OTUs than skin tissue OTUs as shown through the observed OTU number scatter plot and box plot **(Fig. 2A)**. However, the Shannon Index does not show this difference as shown through the Shannon scatter plot and box plot **(Fig. 2B)**, and the heatmap also does not show a difference between the breast and skin OTU abundance **(Fig. 2C)**.

**Figure 2.**
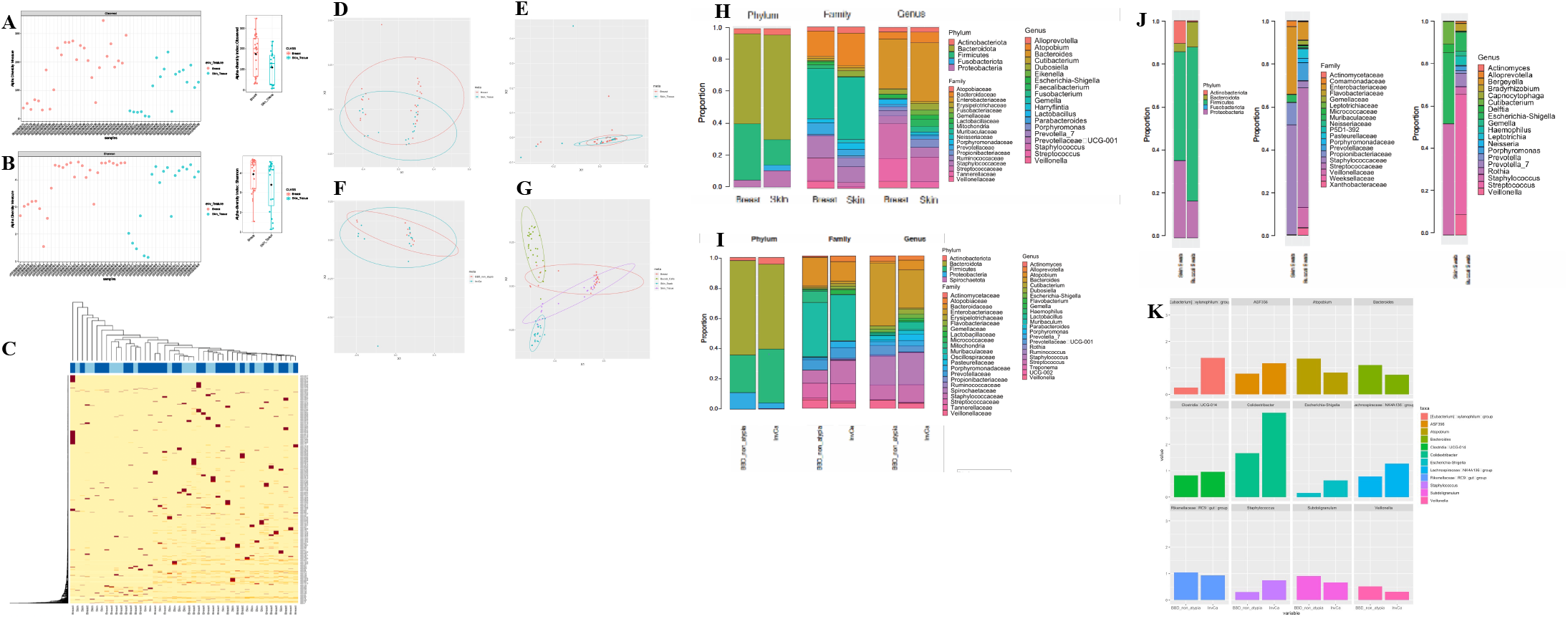
Hieken study re-analysis results. (A) The observed OTU number alpha diversity measure comparing microbiota of breast and skin tissue through a scatter plot and box plot. (B) The Shannon Index alpha diversity measure comparing microbiota of breast and skin tissue through a scatter plot and box plot. (C) Heatmap showing the OTU prevalence of breast (dark blue) and skin (light blue) tissues (columns: samples, rows: OTUs). (D) Unweighted UniFrac distance plot showing clustering of breast (red) and skin (blue) tissues. (E) Weighted UniFrac distance plot showing clustering of breast (red) and skin (blue) tissues. (F) Unweighted UniFrac distance plot showing clustering of breast tissue microbiota between benign (BBD_non_atypia) and cancer (InvCa) states. (G) PCoA plot of unweighted UniFrac distance showing clustering of all 4 tissues where the color scheme is breast tissue (red), skin tissue (purple), skin swab (blue), and buccal swab/cells (green). (H) Proportional abundance barplots generated from our DADA2 analysis of the Hieken data showing taxonomic composition of breast and skin tissue microbiota at the phylum, family, and genus levels. (I) Proportional abundance barplots generated from our DADA2 analysis of the Hieken data showing taxonomic composition of BBD_non_atypia and InvCa samples from breast tissue at the phylum, family, and genus levels. (J) Proportional abundance barplots generated from our DADA2 analysis of the Hieken data showing taxonomic composition of buccal swab and skin swab microbiota at the phylum, family, and genus levels. (K) Differential taxa in breast tissue microbiota of benign (BBD_non_atypia) and malignant disease states based on the linda model.

In the beta diversity analysis, we have used weighted and unweighted UniFrac distances which were calculated through the GUniFrac package in R (Chen et al., 2022) with the OTU table and phylogenetic tree as input. Prior to alpha and beta diversity analyses, the OTU table was rarefied to reduce confounding effects. Beta diversity analysis was performed to compare breast and skin tissue microbiota, where the unweighted UniFrac distance and weighted UniFrac distance did not show a significant difference in microbial community between breast and skin tissue and the MiRKAT p-values for both unweighted and weighted UniFrac were not significant (p > 0.05) **(Fig. 2D; Fig. 2E)**. The microbial community between breast tissue adjacent to invasive cancer disease and breast tissue adjacent to benign disease were also compared using the unweighted and weighted UniFrac distances **(Fig. 2F)**. The unweighted UniFrac distance and weighted UniFrac distance did not show a significant difference in the microbial community between breast tissue adjacent to cancer disease state and breast tissue adjacent to benign disease state and the MiRKAT p-values for both unweighted and weighted UniFrac were not significant (p > 0.05). Additionally, PCoA was performed on the unweighted UniFrac distances and showed that the microbiome of the different tissue types – buccal swab, skin swab, breast tissue, and skin tissue – cluster separately from one another **(Fig. 2G)**. The buccal swab and skin swab microbiota clearly separate from each other and the other two tissues, and the skin and breast tissue microbiota are closer in space but also cluster separate from each other.

In the proportional abundance analysis, we have assessed the taxonomic composition of breast and skin tissue microbiota, breast tissue microbiota in benign, also known as BBD_non_atypia, and invasive cancer, also known as InvCa, disease states, and buccal and skin swab microbiota at phylum, family, and genus levels. The proportional abundance plots include the top 100 sequences in order to clearly show the taxa present at the phylum, family, and genus levels. The breast and skin tissue show similar abundances of major taxa from phyla Actinobacteriota, Bacteroidota, Firmicutes, and Proteobacteria; however, the skin tissue also shows taxa from the phylum Fusobacteriota **(Fig. 2H)**. The breast tissue microbiota in benign and invasive cancer disease states show similar abundances of taxa from phyla Actinobacteriota, Bacteroidota, Firmicutes, and Proteobacteria; however, the invasive cancer disease state shows a low abundance of phylum Spirochaetota **(Fig. 2I)**. The buccal and skin swab show similar abundances of taxa from phyla Actinobacteriota, Bacteroidota, Firmicutes, and Proteobacteria with Fusobacteriota found only in the skin swab **(Fig. 2J)**. At the genus level, there are clearer differences in taxonomic composition between the buccal and skin swab microbiota. The buccal swab microbiota shows a greater abundance of *Veillonella* and *Streptococcus*, whereas the skin swab microbiota shows a greater abundance of *Staphylococcus* and *Escherichia-Shigella*.

In the differential abundance analysis, taxa with prevalence of less than 10% and relative abundance of less than 0.2% were filtered out. In order to identify the differentially abundant taxa, we implemented a linear (lin) model for differential abundance (da) called linda which fits linear regression models on high dimensional data (Zhou et al., 2022) and the linda tool is in the MicrobiomeStat package in R (Zhang et al., 2022). Based on this permutation test, there were twelve significant differentially abundant taxa identified in breast tissue microbiota in benign and invasive cancer disease states. These twelve significant differential taxa were *Eubacterium xylanophilum group, ASF356, Atopobium, Bacteroides, Clostridia UCG-014, Colidextribacter, Escherichia-Shigella, Lachnospiraceae NK4A136 group, Rikenellaceae RC9 gut group, Staphylococcus, Subdoligranulum*, and *Veillonella* where the reported p-values were unadjusted for false discovery correction. The barplots further confirm the abundances of the twelve differential taxa between the benign and malignant disease states in breast tissue **(Fig. 2K)**.

### Chan et al. Re-analysis Results

In the alpha diversity analysis, we have implemented the observed OTU number along with a nonparametric t-test on NS, NAF, and PBS microbiota. The observed OTU number shows the number of OTUs observed for the NS, NAF, and PBS tissues. The t-test assesses whether the microbial diversity is significantly different between the healthy control and cancer samples from NS, NAF, and PBS environments. The p-values are not significant for NS, NAF, and PBS samples **(Fig. 3A; Fig. 3B; Fig. 3C)**.

**Figure 3.**
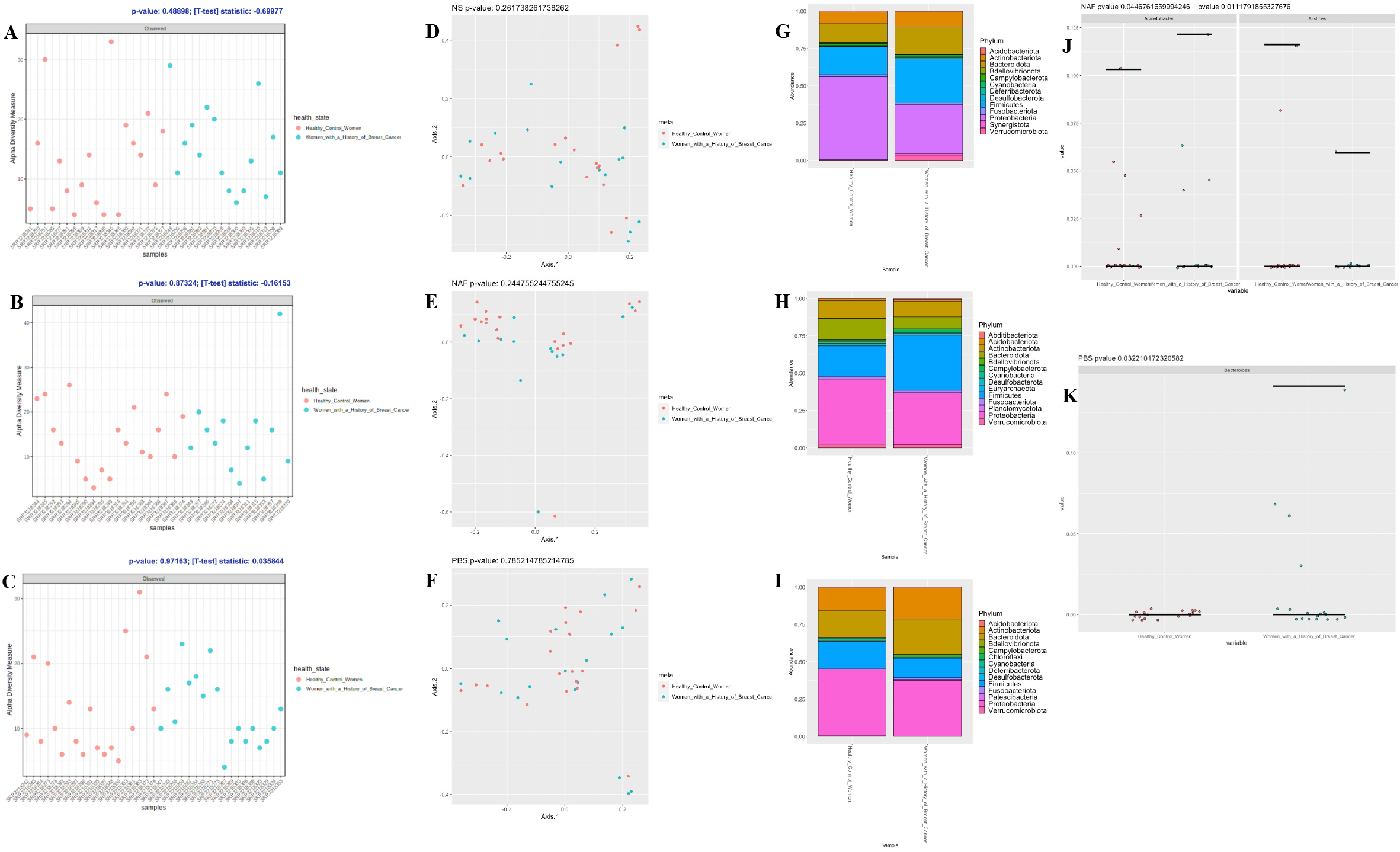
Chan study re-analysis results. (A) NS Observed OTU metric with p-value of 0.489. (B) NAF Observed OTU metric with p-value of 0.873. (C) PBS Observed OTU metric with p-value of 0.972. (D-F) Comparing healthy control women and women with a history of breast cancer across (D) NS, (E) NAF, and (F) PBS samples. (G-I) Comparing proportional abundance of healthy control women and women with a history of breast cancer across (G) NS, (H) NAF, and (I) PBS samples. (J) NAF: Kruskal-Wallis test result of differentially abundant genera, Acinetobacter and Alistipes. (K) PBS: Kruskal-Wallis test result of differentially abundant genus Bacteroides.

In the beta diversity analysis, we have implemented a Bray-Curtis dissimilarity metric and performed PCoA using the rarefied OTU abundances as input where the genus-level OTUs were used in PCoA. The Adonis test was used to test for compositional differences, and the adonis function in the vegan package (Oksanen et al., 2022) from R was used to implement this test. The healthy control and cancer samples from the NS microbiota appear to separate into clusters, but when running the Adonis test there were no significant differences found in the bacterial composition with an unadjusted p-value of 0.262 **(Fig. 3D)**. The healthy control and cancer samples from the NAF microbiota appear to separate into clusters, but when running the Adonis test there were no significant differences found in the bacterial composition with an unadjusted p-value of 0.254 **(Fig. 3E)**. The healthy control and cancer samples from the PBS microbiota appear to separate into clusters, but when running the Adonis test there were no significant differences found in the bacterial composition with an unadjusted p-value of 0.785 **(Fig. 3F)**.

In the proportional abundance analysis, we have assessed the taxonomic composition of NS, NAF, and PBS microbiota at the phylum level. The proportional abundance plots include the top 300 sequences in order to clearly show the taxa present at the phylum level. The NS microbial composition was predominantly comprised of the phyla Proteobacteria, Firmicutes, and Bacteroidota **(Fig. 3G)**. The PBS microbial composition was predominantly comprised of the phyla Proteobacteria, Firmicutes, Bacteroidota, and Actinobacteriota **(Fig. 3I)**. The NAF microbial composition was predominantly comprised of the phyla Proteobacteria and Firmicutes **(Fig. 3H)**.

In the differential abundance analysis, the Kruskal-Wallis test was performed on the healthy and cancer samples from NS, NAF, and PBS OTUs and this test was performed through the kruskal.test function in the stats package (Bolar, 2019) in R. The NS OTUs were not significantly different when comparing the healthy and cancer samples through the Kruskal-Wallis test. There was one PBS OTU identified to be significantly different in relative abundance when comparing the healthy and cancer samples through the Kruskal-Wallis test, and it was *Bacteroides* at the genus level present in only cancer PBS samples with an unadjusted p-value of 0.032 **(Fig. 3K)**. There were two NAF OTUs identified to be significantly different in relative abundance when comparing the healthy and cancer samples through the Kruskal-Wallis test, and they were *Alistipes* and *Acinetobacter* at the genus level **(Fig. 3J)** present in both healthy and cancer NAF samples with an unadjusted p-value of 0.045 for *Acinetobacter* and 0.011 for *Alistipes*.

### Urbaniak et al. Re-analysis Results

In the beta diversity analysis, we performed unsupervised k-means clustering on CLR-transformed data. The clr function in the R package compositions (Boogart et al., 2022) was used to CLR-transform the data and the pam function in the R package cluster (Rousseeuw et al., 2022) was used to perform the unsupervised k-means clustering. There is a clear separation between the breast tumor (BT) cancer samples and the healthy (H) samples as shown in the clusterplot **(Fig. 4A)** where the two components explain 67.18% of the variability.

**Figure 4.**
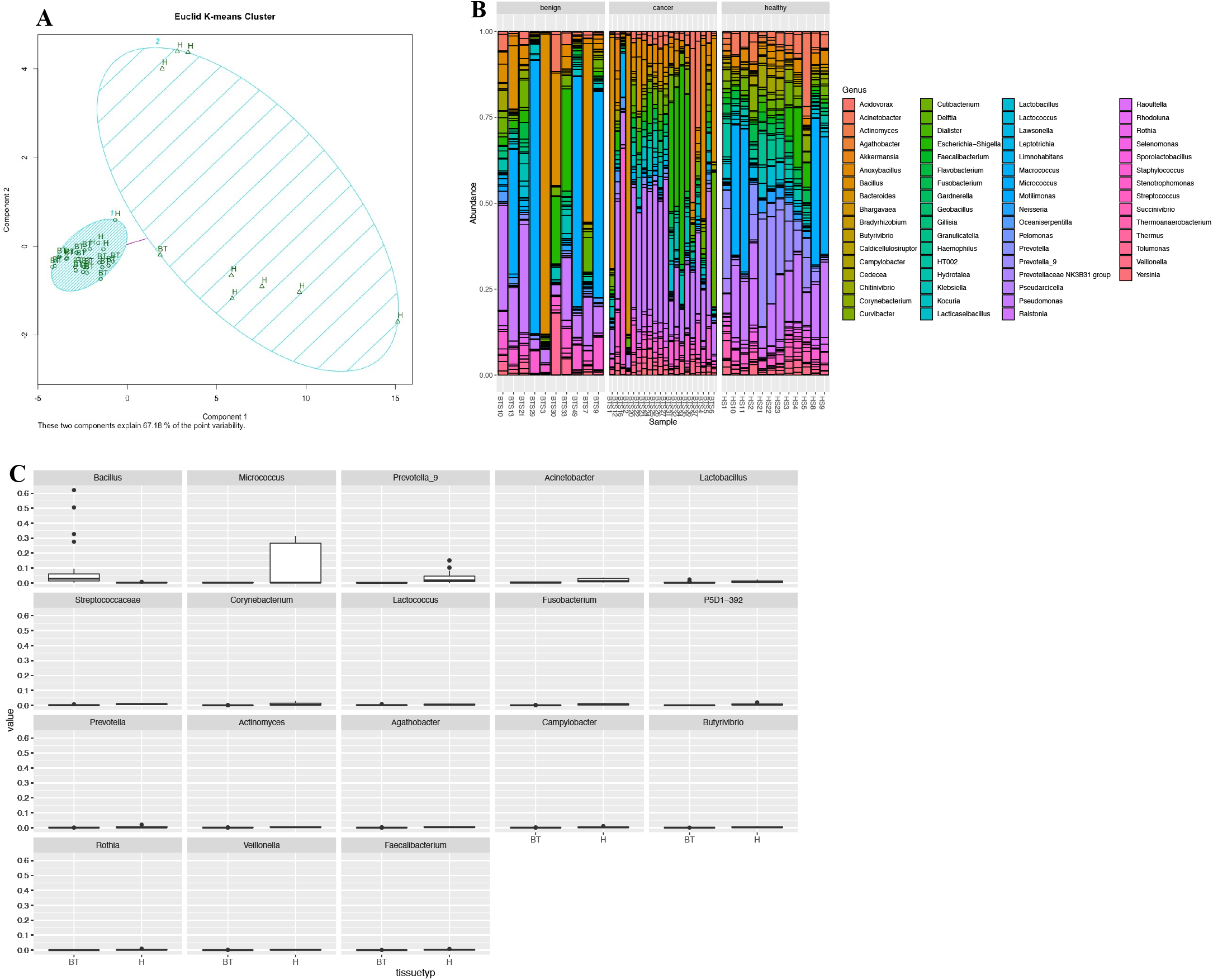
Urbaniak study re-analysis results. (A) K-means clustering plot of CLR-transformed ASV table output from the DADA2 analysis. (B) Proportional abundance plot made using the ASV table output from the DADA2 analysis and taxonomic assignment based on SILVA version 138, with benign (left), cancer (middle), and healthy (right) samples. (C) The 18 statistically significant taxa visualized through boxplots where on x-axis is healthy (right) and breast tumor (left) and y-axis is the abundance.

In the proportional abundance analysis, we assessed the taxonomic composition of healthy, cancer, and benign samples at the genus level. The genus level proportional abundance plot showed a diverse population of bacteria containing 65 genera and top 100 OTUs **(Fig. 4B)**. The proportional abundance plot consists of *Pseudomonas* and *Bacillus* across the benign, cancer, and healthy samples, where *Escherichia-Shigella* was shown to be prevalent in the cancer samples.

In the differential abundance analysis, the ALDEx R package version 2 (Gloor et al., 2022) was used to compare relative abundances of taxa at the genus level of CLR-transformed data. The reported p-values from ALDEx2 are Benjamini-Hochberg corrected p-values of the Wilcoxon rank test. The ALDEx2 output was visualized through boxplots **(Fig. 4C)** and there were 20 statistically significant OTUs and 16 statistically significant taxa of which the following genera were significantly higher in abundance in healthy samples *Acinetobacter, Prevotella_9, Lactobacillus, Corynebacterium, Lactococcus, Fusobacterium, Prevotella, Actinomyces, Agathobacter, Campylobacter, Butyrivibrio, Rothia, Veillonella, Faecalibacterium*, and *Micrococcus*. There was a significantly higher abundance of the following genus in breast tumor cancer samples *Bacillus*.

## Discussion

The individual studies implemented tools, pipelines, and methods that are outdated and are not currently used for bioinformatic analyses, such as IM-TORNADO (Jeraldo, 2016/2020) and the Greengenes reference database (McDonald et al., 2012). While we have observed results that are similar to that of each original study’s results such as when distinguishing between cancer and healthy samples, there are differences present. Our results have identified more taxa than the original studies and have shown that some taxa identified to be significant in the original paper are not found to be significant when re-analyzed with more recent and up-to-date techniques and methods. As mentioned in the introduction, differences in both the bioinformatic pipeline and reference database can contribute to differences in taxonomic assignment and downstream analyses results. The methods implemented in our re-analysis are updated and known to perform better than their older counterparts. The ASV table approach through DADA2 provides greater resolution and lowers false positives (Prodan et al., 2020), and the SILVA database is known to perform better than the Greengenes database (Almeida et al., 2018).

Additionally, the Hieken et al., Chan et al., and Urbaniak et al. studies are seminal papers in the breast cancer microbiome field, and these studies were not able to get a complete and accurate picture of the breast microbiome. Through our re-analysis, we are able to improve upon their results using modern best practices and have discovered new phenomenon that are hitherto unknown and are able to correct their mistaken findings. These findings are important as they will help to elucidate the direction that researchers interested in the breast cancer microbiome should move towards to investigate the appropriate taxa found in the breast microbiome. Therefore, re-analyses of past results and studies are needed to provide the most accurate and reliable results in the scientific community.

The results found from our re-analysis offer new insight into the breast tissue microbial community in healthy, benign, and malignant disease states across different variable regions and patient cohorts. The findings from our re-analysis are able to identify and define more taxa than what was previously reported in each study. There are some overlaps between the findings from our re-analysis, such as *Escherichia-Shigella* is found to be differentially abundant in invasive cancer breast tissue in the Hieken et al. re-analysis and it was also found to be prevalent in the cancer samples in the proportional abundance plot of the Urbaniak et al. re-analysis. Also, *Veillonella* is found to be differentially abundant in benign breast tissue in the Hieken et al. re-analysis and is also found to be differentially abundant in healthy breast tissue in the Urbaniak et al. re-analysis, and, similarly, *Acinetobacter* is found to be differentially abundant in the NAF cancer disease state in the Chan et al. re-analysis and in healthy breast tissue in the Urbaniak et al. re-analysis. There are similarities across the re-analysis results for each study, especially when the tissue and disease states are similar as shown between the Urbaniak et al. re-analysis and Hieken et al. re-analysis, and these similarities may shine light on the microbial communities that could be present in the breast microbiome across disease states.

## Conclusions

In our reevaluation of these three microbiome studies (Chan et al., 2016; Hieken et al., 2016; Urbaniak et al., 2016), we discovered that reanalyses are necessary for every study that studies the microbiome, especially older 16S studies. Reanalyses are important because they provide new insights to the microbiome field and help to assess robusticity of previously published findings by using new and updated tools and databases. In our reevaluation, there were false positives found from the original studies’ results as taxa that were originally identified to be significant were not found to be significant in our reanalyses. Additionally, new taxa were identified when assessing microbial abundance of which were not reported in the original studies’ findings, and multiple taxa were found to be significant between different disease states across the three studies that were not reported in the original studies’ results. Through our reanalyses, we have deduced that every study that studies the microbiome, especially 16S studies, should be continuously reevaluated as the tools and databases develop.

## Availability of data and materials

All data analyzed in this study are obtained from public sources. The data from the Hieken et al. study is available through this accession ID, PRJNA335375, which is stated in their paper as well. The data from the Urbaniak et al. study is available through this accession ID, SRP076038, which is stated in their paper as well. The data from the Chan et al. study is available through this accession ID, PRJNA314877, which is stated in their paper as well.

## Acknowledgements

Thank you to the Hieken et al., Urbaniak et al., and Chan et al. studies for making their data publicly available.

## Ethics declarations

The authors declare no competing interests.

## References

A custom color palette for improving data visualization. (n.d.). Retrieved May 17, 2022, from https://karstenslab.github.io/microshades/

Almeida, A., Mitchell, A. L., Tarkowska, A., & Finn, R. D. (2018). Benchmarking taxonomic assignments based on 16S rRNA gene profiling of the microbiota from commonly sampled environments. GigaScience, 7(5), giy054. https://doi.org/10.1093/gigascience/giy054

Altschul, S. F., Gish, W., Miller, W., Myers, E. W., & Lipman, D. J. (1990). Basic local alignment search tool. Journal of Molecular Biology, 215(3), 403–410. https://doi.org/10.1016/S0022-2836(05)80360-2

American Cancer Society: About Breast Cancer. (n.d.). Key Statistics for Breast Cancer. Retrieved June 21, 2022, from https://www.cancer.org/cancer/breast-cancer/about/how-common-is-breast-cancer.html

Babraham Bioinformatics—FastQC A Quality Control tool for High Throughput Sequence Data. (n.d.). Retrieved May 11, 2022, from https://www.bioinformatics.babraham.ac.uk/projects/fastqc/

Bolar, K. (2019). stat: Interactive Document for Working with Basic Statistical Analysis. https://cran.r-project.org/web/packages/STAT/STAT.pdf

Boogart, K. G. van den, Tolosana-Delgado, R., & Bren, M. (2022). compositions: Compositional Data Analysis. https://cran.r-project.org/web/packages/compositions/compositions.pdf

Burns, M. B., Lynch, J., Starr, T. K., Knights, D., & Blekhman, R. (2015). Virulence genes are a signature of the microbiome in the colorectal tumor microenvironment. Genome Medicine, 7(1), 55. https://doi.org/10.1186/s13073-015-0177-8

Callahan, B. J., McMurdie, P. J., Rosen, M. J., Han, A. W., Johnson, A. J. A., & Holmes, S. P. (2016). DADA2: High-resolution sample inference from Illumina amplicon data. Nature Methods, 13(7), 581–583. https://doi.org/10.1038/nmeth.3869

Caporaso, J. G., Kuczynski, J., Stombaugh, J., Bittinger, K., Bushman, F. D., Costello, E. K., Fierer, N., Peña, A. G., Goodrich, J. K., Gordon, J. I., Huttley, G. A., Kelley, S. T., Knights, D., Koenig, J. E., Ley, R. E., Lozupone, C. A., McDonald, D., Muegge, B. D., Pirrung, M., … Knight, R. (2010). QIIME allows analysis of high-throughput community sequencing data. Nature Methods, 7(5), 335–336. https://doi.org/10.1038/nmeth.f.303

Chan, A. A., Bashir, M., Rivas, M. N., Duvall, K., Sieling, P. A., Pieber, T. R., Vaishampayan, P. A., Love, S. M., & Lee, D. J. (2016). Characterization of the microbiome of nipple aspirate fluid of breast cancer survivors. Scientific Reports, 6(1), 28061. https://doi.org/10.1038/srep28061

Chen, J., Zhang, X., & Yang, L. (2022). GUniFrac: Generalized UniFrac Distances, Distance-Based Multivariate Methods and Feature-Based Univariate Methods for Microbiome Data Analysis. https://cran.r-project.org/web/packages/GUniFrac/GUniFrac.pdf

Cole, J. R., Chai, B., Farris, R. J., Wang, Q., Kulam, S. A., McGarrell, D. M., Garrity, G. M., & Tiedje, J. M. (2005). The Ribosomal Database Project (RDP-II): Sequences and tools for high-throughput rRNA analysis. Nucleic Acids Research, 33(suppl_1), D294–D296. https://doi.org/10.1093/nar/gki038

Computational Biology Core—Brown University. (n.d.). Retrieved May 17, 2022, from https://compbiocore.github.io/metagenomics-workshop/

DADA2 1.16 Pipeline. (n.d.). DADA2 Pipeline Tutorial (1.16). https://benjjneb.github.io/dada2/tutorial.html.

Dhariwal, A., Chong, J., Habib, S., King, I. L., Agellon, L. B., & Xia, J. (2017). MicrobiomeAnalyst: A web-based tool for comprehensive statistical, visual and meta-analysis of microbiome data. Nucleic Acids Research, 45(Web Server issue), W180–W188. https://doi.org/10.1093/nar/gkx295

Edgar, R. C. (2004). MUSCLE: A multiple sequence alignment method with reduced time and space complexity. BMC Bioinformatics, 5(1), 113. https://doi.org/10.1186/1471-2105-5-113

Edgar, R. C. (2010). Search and clustering orders of magnitude faster than BLAST. Bioinformatics, 26(19), 2460–2461. https://doi.org/10.1093/bioinformatics/btq461

Ewels, P., Duncan, A., & Fellows Yates, J. (n.d.). SRA-Explorer. SRA-Explorer. https://sra-explorer.info/

Gloor, G., Fernandes, A., Macklaim, J., Albert, A., Links, M., Quinn, T., Wu, J. R., Wong, R. G., & Lieng, B. (2022). ALDEx2: Analysis Of Differential Abundance Taking Sample Variation Into Account. https://bioconductor.org/packages/release/bioc/manuals/ALDEx2/man/ALDEx2.pdf

Hieken, T. J., Chen, J., Hoskin, T. L., Walther-Antonio, M., Johnson, S., Ramaker, S., Xiao, J., Radisky, D. C., Knutson, K. L., Kalari, K. R., Yao, J. Z., Baddour, L. M., Chia, N., & Degnim, A. C. (2016). The Microbiome of Aseptically Collected Human Breast Tissue in Benign and Malignant Disease. Scientific Reports, 6. https://doi.org/10.1038/srep30751

Jari Oksanen, Gavin L. Simpson, F. Guillaume Blanchet, Roeland Kindt, Pierre Legendre, Peter R. Minchin, R.B. O’Hara, Peter Solymos, M. Henry H. Stevens, Eduard Szoecs, Helene Wagner, Matt Barbour, Michael Bedward, Ben Bolker, Daniel Borcard, Gustavo Carvalho, Michael Chirico, Miquel De Caceres, Sebastien Durand, … James Weedon. (2022). Vegan: Community Ecology Package. https://cran.r-project.org/web/packages/vegan/vegan.pdf

Jeraldo, P. (2020). IM-TORNADO: A pipeline for 16S reads from paired-end libraries [Shell]. https://github.com/pjeraldo/imtornado2 (Original work published 2016)

Kanehisa, M., & Goto, S. (2000). KEGG: Kyoto Encyclopedia of Genes and Genomes. Nucleic Acids Research, 28(1), 27–30.

Langille, M. G. I., Zaneveld, J., Caporaso, J. G., McDonald, D., Knights, D., Reyes, J. A., Clemente, J. C., Burkepile, D. E., Vega Thurber, R. L., Knight, R., Beiko, R. G., & Huttenhower, C. (2013). Predictive functional profiling of microbial communities using 16S rRNA marker gene sequences. Nature Biotechnology, 31(9), 814–821. https://doi.org/10.1038/nbt.2676

Lozupone, C., & Knight, R. (2005). UniFrac: A New Phylogenetic Method for Comparing Microbial Communities. Applied and Environmental Microbiology, 71(12), 8228–8235. https://doi.org/10.1128/AEM.71.12.8228-8235.2005

Lozupone, C., Lladser, M. E., Knights, D., Stombaugh, J., & Knight, R. (2011). UniFrac: An effective distance metric for microbial community comparison. The ISME Journal, 5(2), 169–172. https://doi.org/10.1038/ismej.2010.133

Martin, M. (2011). Cutadapt removes adapter sequences from high-throughput sequencing reads. EMBnet.Journal, 17(1), 10–12. https://doi.org/10.14806/ej.17.1.200

McDonald, D., Price, M. N., Goodrich, J., Nawrocki, E. P., DeSantis, T. Z., Probst, A., Andersen, G. L., Knight, R., & Hugenholtz, P. (2012). An improved Greengenes taxonomy with explicit ranks for ecological and evolutionary analyses of bacteria and archaea. The ISME Journal, 6(3), 610–618. https://doi.org/10.1038/ismej.2011.139

McLaren, M. R., & Callahan, B. J. (2021). Silva 138.1 prokaryotic SSU taxonomic training data formatted for DADA2 [Data set]. Zenodo. https://doi.org/10.5281/zenodo.4587955

Mira-Pascual, L., Cabrera-Rubio, R., Ocon, S., Costales, P., Parra, A., Suarez, A., Moris, F., Rodrigo, L., Mira, A., & Collado, M. C. (2015). Microbial mucosal colonic shifts associated with the development of colorectal cancer reveal the presence of different bacterial and archaeal biomarkers. Journal of Gastroenterology, 50(2), 167–179. https://doi.org/10.1007/s00535-014-0963-x

Price, M. N., Dehal, P. S., & Arkin, A. P. (2009). FastTree: Computing Large Minimum Evolution Trees with Profiles instead of a Distance Matrix. Molecular Biology and Evolution, 26(7), 1641–1650. https://doi.org/10.1093/molbev/msp077

Prodan, A., Tremaroli, V., Brolin, H., Zwinderman, A. H., Nieuwdorp, M., & Levin, E. (2020). Comparing bioinformatic pipelines for microbial 16S rRNA amplicon sequencing. PLOS ONE, 15(1), e0227434. https://doi.org/10.1371/journal.pone.0227434

Pruesse, E., Quast, C., Knittel, K., Fuchs, B. M., Ludwig, W., Peplies, J., & Glöckner, F. O. (2007). SILVA: A comprehensive online resource for quality checked and aligned ribosomal RNA sequence data compatible with ARB. Nucleic Acids Research, 35(21), 7188–7196. https://doi.org/10.1093/nar/gkm864

Rousseeuw, P., Struyf, A., & Hubert, M. (2022). cluster: Methods for Cluster analysis. https://cran.r-project.org/web/packages/cluster/cluster.pdf

Schliep, K., Paradis, E., Martins, L. de O., Potts, A., White, T. W., Stachniss, C., Kendall, M., Halabi, K., Bilderbeek, R., Winchell, K., Revell, L., Gilchrist, M., Beaulieu, J., O’Meara, B., & Qu, L. (2021). phangorn: Phylogenetic Reconstruction and Analysis. https://cran.r-project.org/web/packages/phangorn/phangorn.pdf

Schloss, P. D., Westcott, S. L., Ryabin, T., Hall, J. R., Hartmann, M., Hollister, E. B., Lesniewski, R. A., Oakley, B. B., Parks, D. H., Robinson, C. J., Sahl, J. W., Stres, B., Thallinger, G. G., Horn, D. J. V., & Weber, C. F. (2009). Introducing mothur: Open-Source, Platform-Independent, Community-Supported Software for Describing and Comparing Microbial Communities. Applied and Environmental Microbiology, 75(23), 7537–7541. https://doi.org/10.1128/AEM.01541-09

Sequence Read Archive (SRA). (n.d.). National Center for Biotechnology Information, National Library of Medicine (US). https://www.ncbi.nlm.nih.gov/sra/

Urbaniak, C., Gloor, G. B., Brackstone, M., Scott, L., Tangney, M., & Reid, G. (2016). The Microbiota of Breast Tissue and Its Association with Breast Cancer. Applied and Environmental Microbiology, 82(16), 5039–5048. https://doi.org/10.1128/AEM.01235-16

Xuan, C., Shamonki, J. M., Chung, A., DiNome, M. L., Chung, M., Sieling, P. A., & Lee, D. J. (2014). Microbial Dysbiosis Is Associated with Human Breast Cancer. PLOS ONE, 9(1), e83744. https://doi.org/10.1371/journal.pone.0083744

Zhang, X., Chen, J., & Zhou, H. (2022). MicrobiomeStat: Statistical Methods for Microbiome Compositional Data. https://cran.r-project.org/web/packages/MicrobiomeStat/MicrobiomeStat.pdf

Zhao, N., Chen, J., Carroll, I. M., Ringel-Kulka, T., Epstein, M. P., Zhou, H., Zhou, J. J., Ringel, Y., Li, H., & Wu, M. C. (2015). Testing in Microbiome-Profiling Studies with MiRKAT, the Microbiome Regression-Based Kernel Association Test. American Journal of Human Genetics, 96(5), 797–807. https://doi.org/10.1016/j.ajhg.2015.04.003

Zhou, H., He, K., Chen, J., & Zhang, X. (2022). LinDA: Linear models for differential abundance analysis of microbiome compositional data. Genome Biology, 23(1), 95. https://doi.org/10.1186/s13059-022-02655-5

